# AdapToR: Adaptive Topological Regression for quantitative structure-activity relationship modeling

**DOI:** 10.1101/2025.04.02.646801

**Authors:** Yixiang Mao, Souparno Ghosh, Ranadip Pal

## Abstract

Quantitative structure-activity relationship (QSAR) modeling has become a critical tool in drug design. Recently proposed Topological Regression (TR), a computationally efficient and highly interpretable QSAR model that maps distances in the chemical domain to distances in the activity domain, has shown predictive performance comparable to state-of-the-art deep learning-based models. However, TR’s dependence on simple random sampling-based anchor selection and utilization of radial basis function for response reconstruction constrain its interpretability and predictive capacity. To address these limitations, we propose Adaptive Topological Regression (AdapToR) with adaptive anchor selection and optimization-based reconstruction. We evaluated AdapToR on the NCI60 GI50 dataset, which consists of over 50,000 drug responses across 60 human cancer cell lines, and compared its performance to Transformer CNN, Graph Transformer, TR, and other baseline models. The results demonstrate that AdapToR outperforms competing QSAR models for drug response prediction with significantly lower computational cost and greater interpretability as compared to deep learning-based models.

## 1 Introduction

Quantitative structure-activity relationship (QSAR) modeling predicts the biological activities of chemical compounds from their molecular structures. This modeling technique has become an essential tool to accelerate the drug discovery process, saving time, costs, and resources [1–4]. It is widely used in several phases of drug discovery, including virtual screening for hits identification, hits-to-lead optimization, and lead optimization [1–3]. In virtual screening, QSAR models can help to identify promising compounds from large chemical libraries, reducing the number of compounds for synthesis and assays [1, 2]. In hits-to-lead and lead optimization, QSAR modeling can guide the multi-parameter optimization process by elucidating connections between chemical structures and their biological activities, such as potency, selectivity, and pharmacokinetic parameters [1].

The majority of QSAR models are supervised machine learning (ML) models and can be broadly categorized into feature-based and similarity-based approaches. In feature-based approaches, molecular structures are first transformed into machine-comprehensible features. Common feature types include (a) vectors, such as molecular descriptors and fingerprints [5–7], (b) graphs, and (c) strings, such as Simplified Molecular Input Line Entry System (SMILES). These features are then used to train predictive ML models. In recent decades, there has been a shift from traditional shallow learners, such as linear regression (LR), Support Vector Machines (SVM), and Random Forest (RF) [8–10], towards more sophisticated deep learning (DL)-based QSAR models [11–17]. One group of DL methods utilizes large language models with SMILES strings as input [11, 12]. For example, the Transformer-Convolutional Neural Network (TCNN) model proposed in [12] combines a Transformer encoder with a Text-CNN, exhibiting superior performance on both regression and classification tasks across multiple datasets. Another group of DL models employs Graph Convolutional Networks to learn features from molecular graphs [13, 15–17]. These models are commonly designed to predict cancer drug responses with additional cell line features learned by other DL networks. While these models have demonstrated superior performance on mixed tests (where the test set was randomly picked from all possible drug-cell line pairs), their performance degrades in drug-blind settings (i.e., predicting unseen drugs) [13, 17].

Despite achieving better predictive performance compared to traditional shallow learners, DL-based QSAR models suffer from high computational complexity and lack of interpretability. Recall that model interpretability is different from model explainability. Following the definitions proposed in [18], we view an interpretable model to be one whose construction is inherently interpretable, whereas explainable ML tries to provide post hoc explanations for existing black box models. As an example of explainable QSAR models, [12] used Layer-wise Relevance Propagation algorithm [19, 20] to assign importance scores to input features based on their contribution to the final prediction. Although these scores indicate the relative influence of features on the final prediction, they do not reveal how the model explicitly utilizes these features to obtain the final prediction. In contrast, white box models, such as LR and Decision Trees, are inherently interpretable. Their decision-making processes can be directly examined by analyzing model weights (in LR) or decision nodes (in Decision Trees) [21]. While interpretability may not be critical for hits identification, it can be essential for hits-to-lead and lead optimization, where QSAR models are responsible for linking chemical structures to biological activities.

Similarity-based methods, such as *K* − Nearest Neighbor (KNN) and kernel regression [22], offer intuitive interpretability, as their predictions are typically computed as a weighted sum of the target values of training samples, with weights determined by similarity in the input space. Recently, [23] introduced a novel similarity-based regression approach called Topological Regression (TR), which is statistically robust, computationally efficient, and interpretable. TR builds linear models that use distances in the structure (input) space to predict distances in the response space. These distances are calculated between samples and anchor points that are randomly selected from training samples. When testing, the estimated response distances between a test sample and the anchor points are converted to weights through a Radial Basis Function (RBF). The final prediction is then reconstructed as a weighted sum of the responses of the anchor points. When evaluated on ChEMBL datasets [24, 25], the predictive performance of TR was comparable to that of TCNN, but at a significantly lower computational cost and greater interpretability. In addition, TR outperformed other competing models, including RF, metric learning kernel regression, and ChemProp [26].

Although TR had demonstrated promising predictive capacity and great interpretability, the performance of this framework could be improved by optimizing the process of anchor point selection and response reconstruction methodologies. First, TR selects the anchor points via simple random sampling, resulting in uncertainty in model performance. To mitigate the effect of random anchor selection, [23] proposed Ensemble TR that averages predictions across multiple models trained with different sets of anchor points. While the ensemble approach achieves better predictive performance, it sacrifices computational efficiency and model interpretability. In this work, we propose an adaptive anchor selection strategy to address the uncertainty and to improve model performance. Since this adaptive strategy produces a single set of anchor points, it is more interpretable and time-efficient than the ensemble approach. Furthermore, in the original TR, the RBF-based response reconstruction overemphasized small distance estimates in the response space, while disregarding large distance estimates. However, large distance estimates can also carry meaningful information. For example, if a test sample has a large estimated response distance to an anchor with a low response value, it could indicate a high response value for that test sample. In addition, the RBF-based reconstruction is not optimized in any form. To address these weaknesses, we propose an optimization-based response reconstruction approach that can utilize information across all distance estimations and is optimized under the stated loss criterion. In addition to these innovative approaches, we implemented several other modifications to stabilize model behavior and reduce computation time. We refer to this adaptive version of TR as **Adap**tive **To**pological **R**egression (**AdapToR**).

In [23], TR is evaluated on ChEMBL datasets with small to moderate sample sizes (that range from 100 to 7890 with a median sample size of 677). This evaluation may favor simpler models like TR over DL models that typically require a large number of training samples for optimal performance. In this work, we evaluated TR and proposed AdapToR on the NCI60 GI50 dataset [27] that consists of over 50,000 drug responses across 60 human cancer cell lines. To the best of our knowledge, it is one of the largest real-world datasets for cancer drug response prediction. For comparison, we implemented and evaluated multiple baseline models, including LR, RF, and KNN, as well as the state-of-the-art DL models, including TCNN [12] and Graph Transformer (GraTrans) [17]. Our results show that AdapToR outperforms all competing models while being considerably more time-efficient compared to TR and the DL models. Furthermore, we present an illustrative example to demonstrate the interpretability of a trained AdapToR model and discuss the insights that can be derived from it.

The paper is organized as follows. Section 2 provides a short description of the data and experiments along with the results. Section 3 provides a discussion on the framework. Section 4 provides detailed descriptions of the methods used in the manuscript. The data and code availability are included in Section 7.

## 2 Results

### 2.1 Data and experiment description

#### 2.1.1 NCI60 GI50 dataset

The NCI60 GI50 dataset assesses the anticancer activity of over 50,000 compounds across 60 human cancer cell lines. Drug responses are measured as GI50 values (in molar units, M), representing the concentration required to inhibit 50% of cell proliferation [27]. We transformed the GI50 values into their negative logarithmic form, referred to as NLOGGI50 (NLOGGI50= − log_10_(GI50)), and used that as response values. Higher NLOGGI50 values indicate stronger inhibitory effects.

The drugs’ unique NSC# was used to obtain their SMILES through PubChem. Drugs that do not have valid SMILES strings were excluded. The molecular fingerprints and graphs are then generated based on their SMILES. For fingerprints, we generated Extended-Connectivity Fingerprints with a radius of 2 (ECFP4) using RDKit [28], and MinHash fingerprint with a radius of 3 (MHFP6) [6]. The number of bits was set to 2048 for both fingerprints. ECFP4 was originally used in TR, and MHFP6 has been shown to outperform ECFP4 in nearest neighbor searches [6]. Our empirical results (see in Fig. 3) indicate that using MHFP6 enhances the performance of TR. Molecular graphs were generated following the procedure described in [17] and used as inputs to the GraTrans model.

We excluded the cell line MDA-MB-468, which has less than 10,000 drugs. Thus, our final dataset consisted of 59 cell lines and 51,312 unique drug compounds. Table S1 shows the number of drugs for each cell line.

#### 2.1.2 Comparison procedure

Our goal is to predict drug response for each cell line in a drug-blind setting. The predictive performance of each candidate model was assessed using 5-fold cross-validation (CV), with each fold consisting of 80% training and 20% testing data. The separation of training and testing sets is independent for each cell line. We evaluated the candidate models using normalized root mean square error (NRMSE), Spearman’s rank correlation coefficient (*ρ*), Pearson correlation coefficient (PCC, *r*), and bias in predictions (bias). NRMSE compares the predictive residuals of a trained model to the prediction error obtained from a null model (recall, a null model uses the mean of the training responses as the point prediction for all test samples). Spearman’s *ρ* assesses the monotonic relationship between the observed and predicted responses. It will be close to unity if the predicted and observed responses have similar ranks. PCC, on the other hand, measures the linear relationships between the predicted and observed responses. The bias in prediction is defined as the slope of the best-fit line through the residuals (prediction error) as a function of the observed responses. It evaluates whether the residuals are systematically related to the observed responses. Empirically, if we observe the residuals to be randomly distributed about 0, the bias value will tend to be zero, indicating an unbiased prediction. Denote the observed and predicted responses in the test set as *y*_*i*_ and *ŷ*_*i*_, *i* = 1, 2, …, *n*, respectively. Define the corresponding mean values as 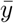 and 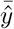, the corresponding estimated standard deviation values as 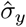 and 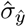, and the difference between the ranks of the observed and predicted responses as *v*_*i*_, *i* = 1, 2, …, *n*, the performance metrics are calculated as follows:

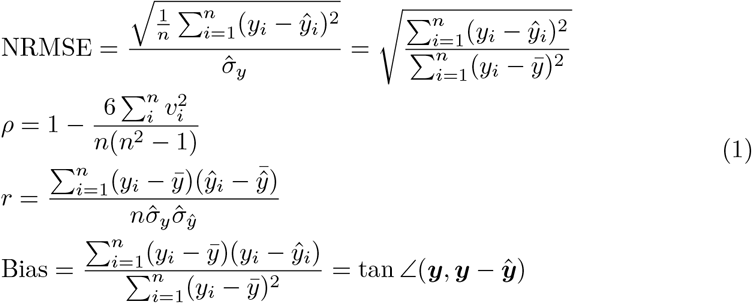

### 2.1.3 GPU and CPU systems

For a fair comparison and to optimize hardware performance for each model, we trained and tested the DL models on a GPU system equipped with an NVIDIA Tesla V100 GPU and an Intel Xeon Cascade Lake 6248 CPU (2.5 GHz, 20 cores, 40 threads). Other models were trained on a CPU system featuring an AMD EPYC ROME 7702 CPU (2.0 GHz, 64 cores, 128 threads).

### 2.2 Model performance comparison

#### 2.2.1 NCI60 GI50 datasets

The average NRMSE, Spearman’s *ρ*, PCC, and bias for each model over 59 cell lines and 5-fold CV splits are listed in Table 1. Fig. 1 shows the boxplots of the performance metrics. As shown in the table and figure, AdapToR achieves the best performance, exhibiting the lowest NRMSE and bias, as well as the highest Spearman’s *ρ* and PCC. TCNN with data augmentation (TCNN-Aug) demonstrates performance that is comparable, though slightly inferior, to AdapToR. Following these two models, TR, TCNN, GraTrans, RF, and KNN show moderate performance, whereas LR performs the worst. To reveal a more fine-grained picture, Fig. 2 shows scatter plots of the NRMSE values of AdapToR versus (a) TCNN-Aug and (b) TR across 59 cell lines and 5 CV splits (a total of 295 data splits). As shown in Fig. 2 (a), for 289 out of the 295 splits (98%), AdapToR achieves less NRMSE than TCNN-Aug. In Fig. 2 (b), AdapToR shows less NRMSE than TR in all data splits.

**Table 1:**
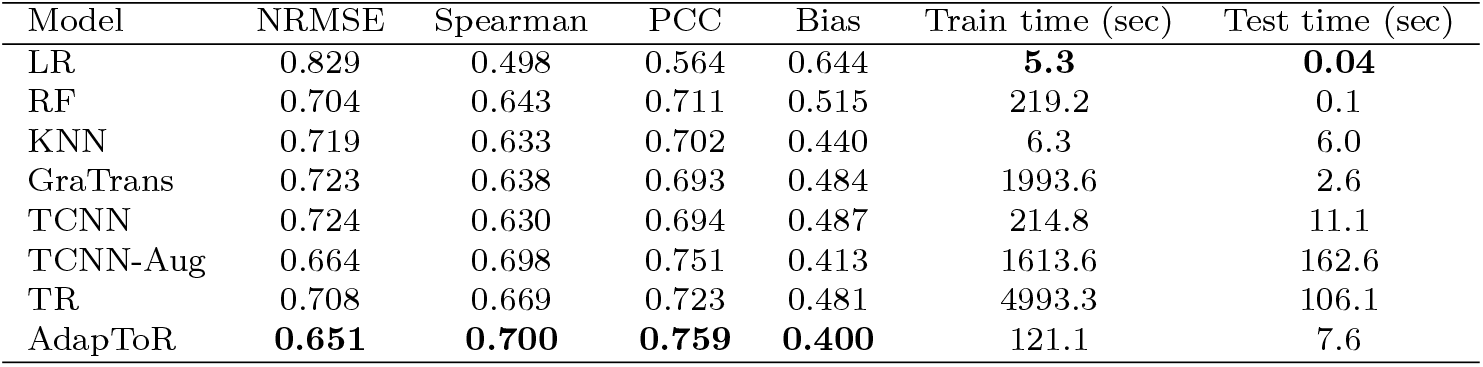
Model performance averaged over 59 cell lines of the NCI60 GI50 dataset and 5-fold cross-validation splits.

| Model | NRMSE | Spearman | PCC | Bias | Train time (sec) | Test time (sec) |
| --- | --- | --- | --- | --- | --- | --- |
| LR | 0.829 | 0.498 | 0.564 | 0.644 | **5.3** | **0.04** |
| RF | 0.704 | 0.643 | 0.711 | 0.515 | 219.2 | 0.1 |
| KNN | 0.719 | 0.633 | 0.702 | 0.440 | 6.3 | 6.0 |
| GraTrans | 0.723 | 0.638 | 0.693 | 0.484 | 1993.6 | 2.6 |
| TCNN | 0.724 | 0.630 | 0.694 | 0.487 | 214.8 | 11.1 |
| TCNN-Aug | 0.664 | 0.698 | 0.751 | 0.413 | 1613.6 | 162.6 |
| TR | 0.708 | 0.669 | 0.723 | 0.481 | 4993.3 | 106.1 |
| AdapToR | **0.651** | **0.700** | **0.759** | **0.400** | 121.1 | 7.6 |

**Fig. 1:**
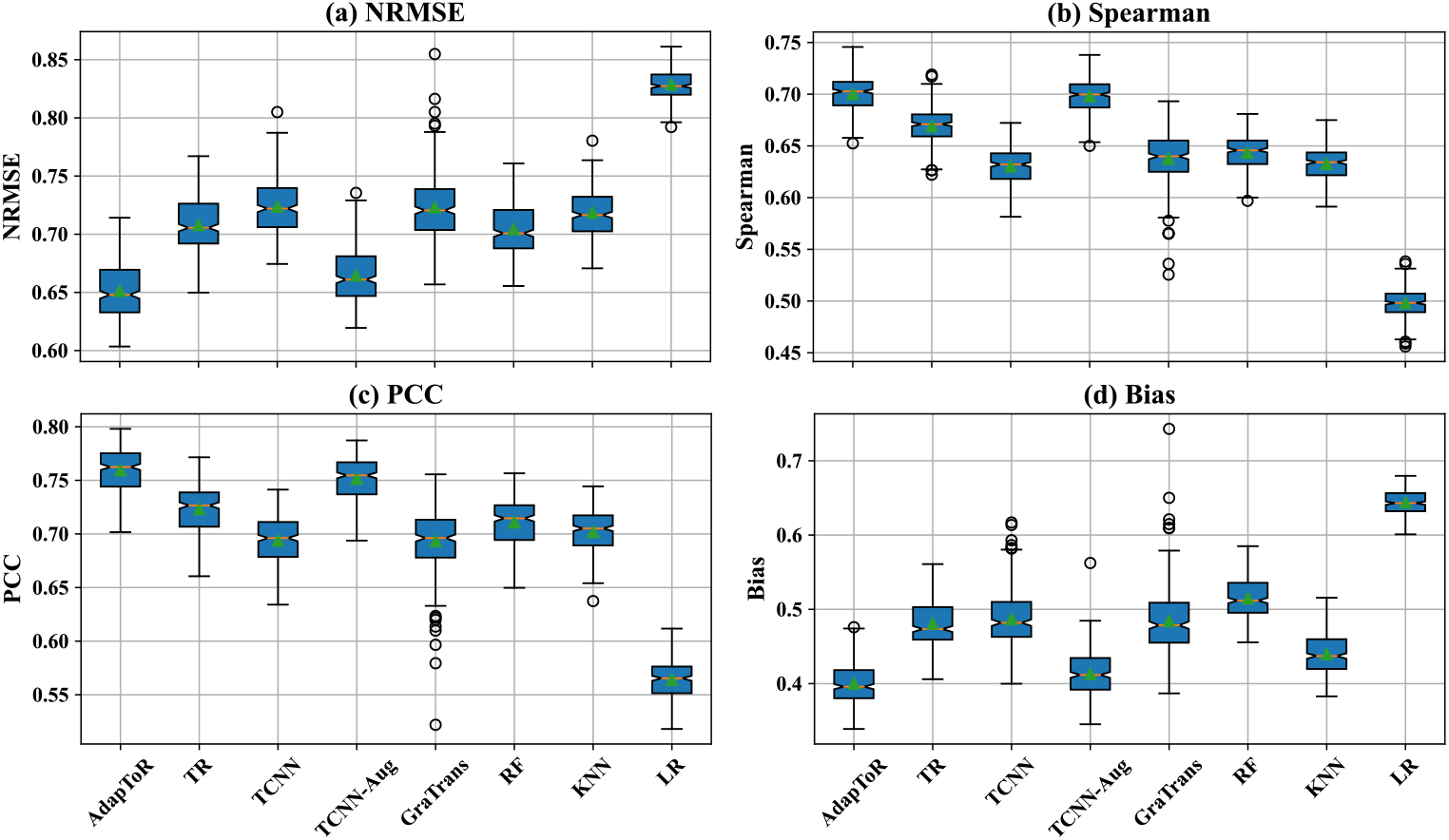
Boxplots showing the (a) NRMSE, (b) Spearman’s rank correlation coefficient, (c) Pearson correlation coefficient, and (d) bias of different models, including AdapToR, TR, TCNN, TCNN-Aug, GraTrans, RF, KNN, and LR. The orange lines and green triangles represent the medians and the means, respectively. The notches indicate the 95% confidence interval of the medians.

**Fig. 2:**
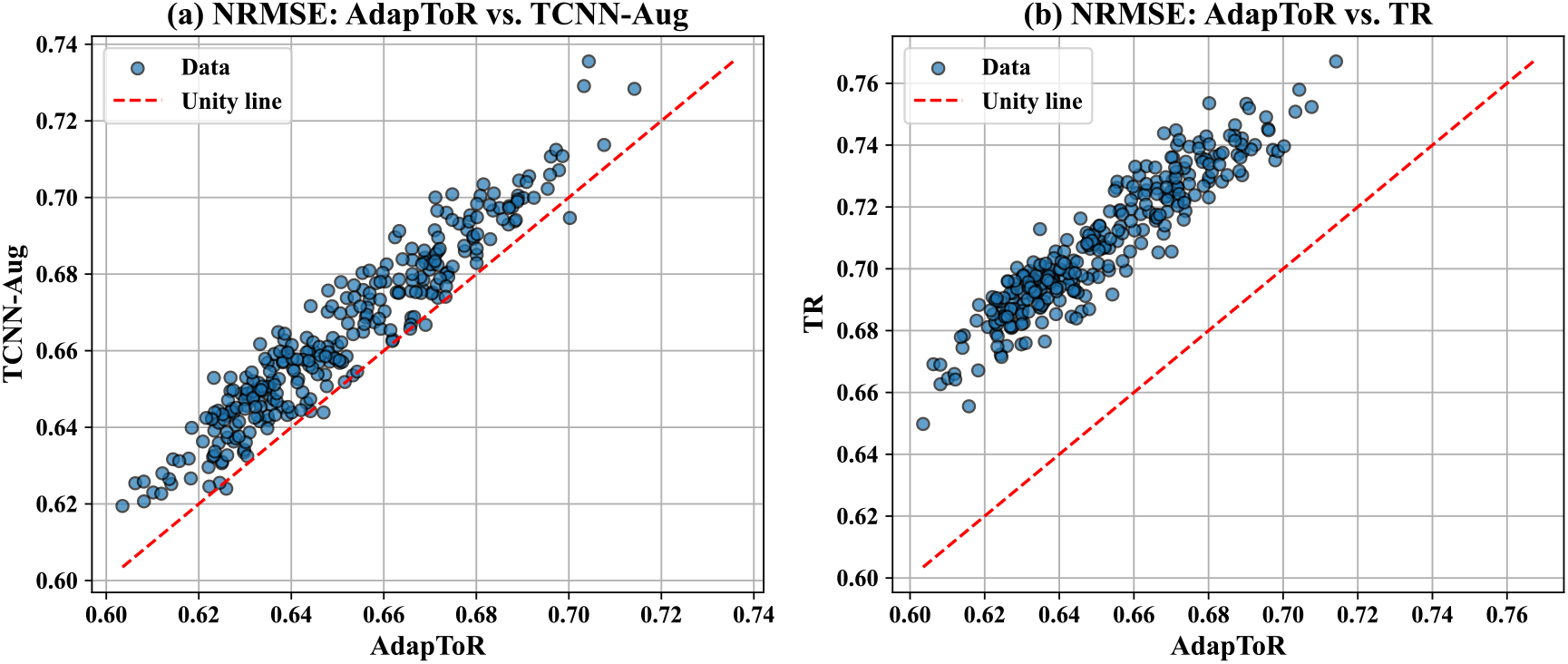
Scatter plots comparing the NRMSE values of AdapToR versus (a) TCNN-Aug and (b) TR across 59 cell lines and 5 CV splits (a total of 295 data splits). The red dashed line represents the unity line.

In addition to the performance metrics, we also compared the time complexity of different models. Table 1 provides the average training and testing times for each model. AdapToR shows moderate training and testing times of 121.1 and 7.6 sec, respectively. Its training time is only longer than LR (5.3 sec) and KNN (6.3 sec) and is significantly lower than TR (4993.3 sec), GraTrans (1993.6 sec), and TCNN-Aug (1613.6 sec). The testing time of AdapToR (7.6 sec) is considerably shorter than TCNN-Aug (162.6 sec) and TR (106.1 sec) and is slightly longer than the baseline models and GraTrans (ranging from 0.04 to 5.9 sec). TR exhibits the longest training time because solving Eq. 4 becomes numerically unstable in the presence of multi-collinearity. TCNN-Aug demonstrates the longest testing time because it generates, evaluates, and averages predictions for multiple augmented samples per test instance. Taken together, when evaluated on the NCI60 GI50 dataset, AdapToR outperforms competing models in all performance metrics (NRMSE, Spearman’s *ρ*, PCC, and bias) and demonstrates an order of magnitude less training and testing times as compared to TCNN-Aug.

#### 2.2.2 ChEMBL datasets

As supplementary results, we trained and evaluated AdapToR on 530 ChEMBL datasets. Detailed descriptions of dataset selection and data processing are available in [23]. Table S2 summarizes the average NRMSE, Spearman’s *ρ*, and training and testing times for AdapToR, TR, Ensemble TR, TCNN, and TCNN-Aug over 530 ChEMBL datasets and 5-fold CV splits. Consistent with previous findings, AdapToR achieves the best performance at much lower training and testing times compared to Ensemble TR, TCNN, and TCNN-Aug.

### 2.3 Ablation analysis of AdapToR

Compared to TR, AdapToR achieves a 41-fold reduction in training time, a 14-fold reduction in testing time, 8.0% and 16.8% reductions in NRMSE and bias, respectively, and an approximate 5% increase in Spearman’s *ρ* and PCC. To better understand the implications of each modification introduced in AdapToR, we performed an ablation analysis of AdapToR.

As shown in Fig. 7 and described in detail in Section 4, AdapToR uses MHFP6 to calculate the structure distances and adds four key model features to the vanilla TR model. These additional features include (1) training a Ridge regression model (denoted as L2) to map structure distances to response distances, (2) selecting 10 response anchors with *k*−means clustering (denoted as Kmeans), (3) novel adaptive anchor selection in the structure space, and (4) novel optimization-based response reconstruction.

Starting with vanilla TR, we incrementally added one modification at a time, resulting in six different models: TR, TR (L2), TR (L2, MHFP6), TR (L2, MHFP6, Kmeans), AdapToR (RBF), and AdapToR. AdapToR (RBF) represents the AdapToR model with optimization-based reconstruction replaced by RBF-based reconstruction. Fig. 3 shows boxplots of the performance metrics, and Table 2 lists the corresponding average values.

**Table 2:**
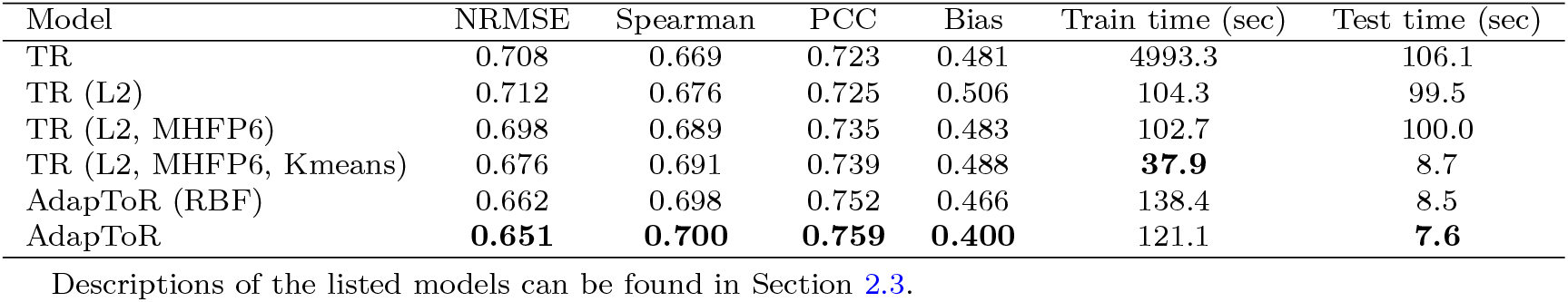
Ablation analysis of AdapToR. Performance of different TR configurations averaged over 59 cell lines of the NCI60 GI50 dataset and 5-fold cross-validation splits

Regarding the performance metrics, all modifications, except for the use of Ridge regression, had a positive impact on the performance metrics, reducing the NRMSE and bias while increasing Spearman’s *ρ* and PCC. Specifically, NRMSE steadily decreased from TR (L2) to AdapToR. For Spearman’s *ρ* and PCC, major improvements are observed after incorporating MHFP6 and adaptive anchor selection. For bias, the largest reduction was attributed to optimization-based reconstruction, followed by adaptive anchor selection.

Regarding time complexity, using Ridge regression leads to a significant reduction in training time. Additionally, selecting 10 response anchors with *k*− means results in shorter training and testing times. Incorporating adaptive anchor selection, however, increases training time because additional time is required to evaluate training samples and train intermediate models during the anchor selection process. Optimization-based reconstruction slightly reduces both training and testing times.

#### 2.3.1 Performance of ensemble and stacking TR

In this section, we assess the performance of AdapToR when replacing adaptive anchor selection with the ensemble approach proposed in [23]. The resulting model is denoted as Ensemble TR (enhanced). The ensemble approach randomly selects 30% to 90% of the training samples as structure anchors to train fifteen TR models, where the percentages are sampled from a Gaussian distribution (mean = 0.6, std = 0.3). The final prediction is obtained by averaging the predictions from these models. In addition to the ensemble approach, we also evaluated the performance of stacking the predictions with a linear model, where 10% of the training samples were randomly selected as the validation set to estimate stacking weights. The resulting model is denoted as Stack TR (enhanced). Moreover, we explored the ensemble and stack approaches within the adaptive anchor selection framework. We averaged or stacked the predictions from the intermediate and final models trained in adaptive anchor selection, leading to AdapToR (ensemble) and AdapToR (stack), respectively.

The performance of these models is summarized in Table 3. As shown in the table, Ensemble TR (enhanced) and Stack TR (enhanced) exhibit lower performance and longer training and testing times compared to AdapToR. Furthermore, averaging or stacking predictions from models trained in adaptive anchor selection does not improve model performance and instead results in increased testing times.

**Table 3:** Performance of ensemble and stacking TR models averaged over 59 cell lines of the NCI60 GI50 dataset and 5-fold cross-validation splits.

| Model | NRMSE | Spearman | PCC | Bias | Train time (sec) | Test time (sec) |
| --- | --- | --- | --- | --- | --- | --- |
| Ensemble TR (enhanced) | 0.662 | 0.695 | 0.750 | 0.418 | 701.7 | 123.4 |
| Stack TR (enhanced) | 0.670 | 0.689 | 0.743 | 0.428 | 582.8 | 123.5 |
| AdapToR (ensemble) | **0.652** | **0.700** | **0.759** | **0.399** | 137.1 | 33.0 |
| AdapToR (stack) | 0.659 | 0.694 | 0.753 | 0.409 | 136.7 | 32.7 |
| AdapToR | **0.651** | **0.700** | **0.759** | **0.400** | **121.1** | **7.6** |

Descriptions of the listed models can be found in Section 2.3.1.

#### 2.3.2 Comparing random and adaptive anchor selection

To better understand how adaptive anchor selection works, Fig. 4 compares the absolute prediction error of AdapToR models trained with random and adaptive anchor selection on 500 samples randomly picked from the HCC2998 cell line. Fig. 4 (a) shows the ground truth response values. Figs. 4 (b) and (c) illustrate the absolute prediction error for the first and second models trained in the adaptive anchor selection process, respectively. The red dots in those figures represent the 75 (15%) randomly selected structure anchors used to train the first model, denoted as ***S***_1_. The orange stars in Fig. 4 (c) indicate the 75 samples not in ***S***_1_ that exhibited the highest absolute prediction error from the first model, denoted as 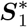. The combination of the red dots and orange stars was used to train the second model. For comparison, Fig. 4 (d) shows the absolute prediction error for the second model when samples in 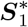 were randomly selected from samples not in ***S***_1_.

**Fig. 3:**
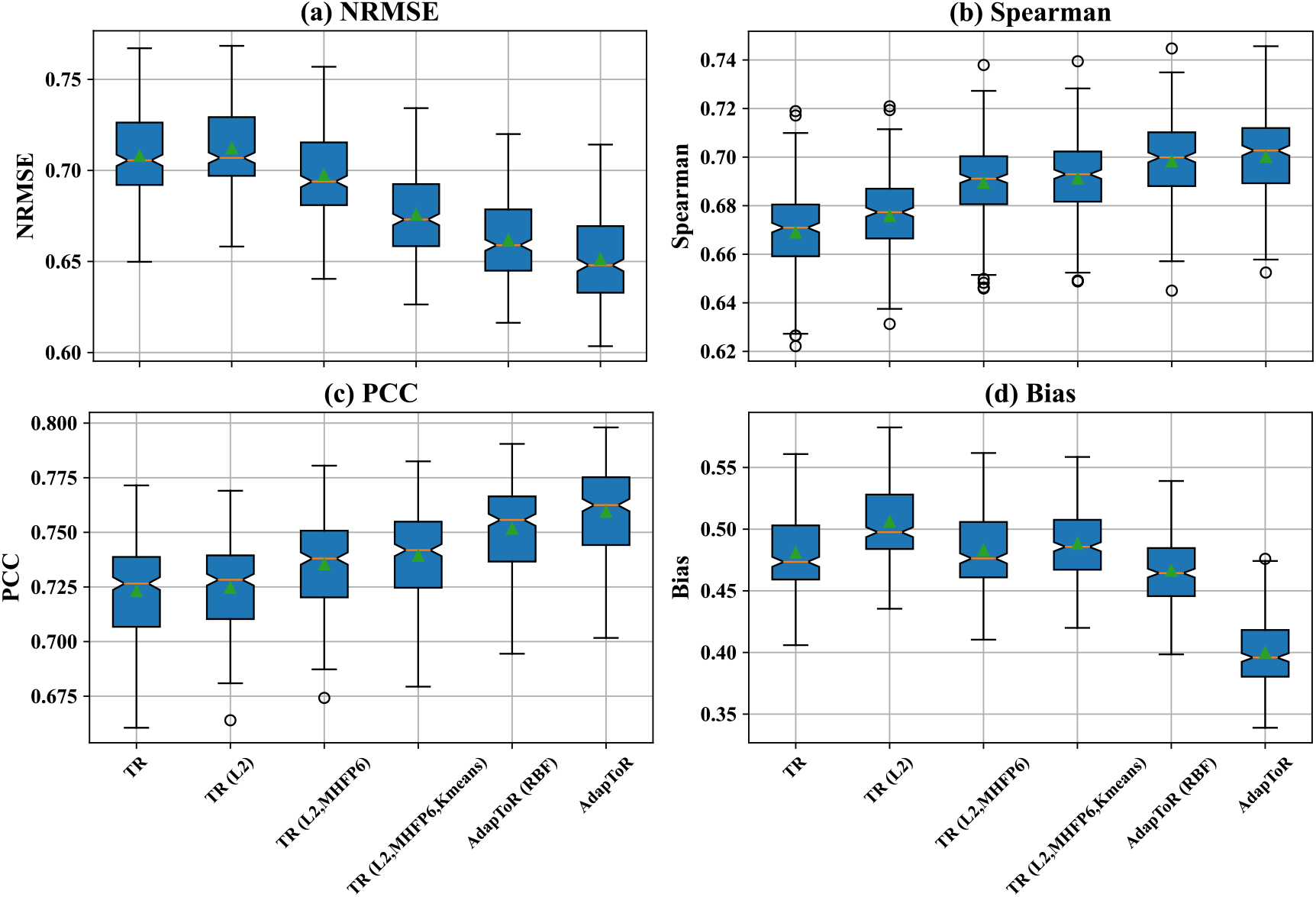
Ablation analysis of AdapToR. Boxplots show the (a) NRMSE, (b) Spearman’s rank correlation coefficient, (c) Pearson correlation coefficient, and (d) bias of different TR models, including TR, TR (L2), TR (L2, MHFP6), TR (L2, MHFP6, Kmeans), AdapToR (RBF) and AdapToR. AdapToR (RBF) represents the AdapToR model with optimization-based reconstruction replaced by RBF-based reconstruction. The orange lines and green triangles represent the medians and the means, respectively. The notches indicate the 95% confidence interval of the medians.

**Fig. 4:**
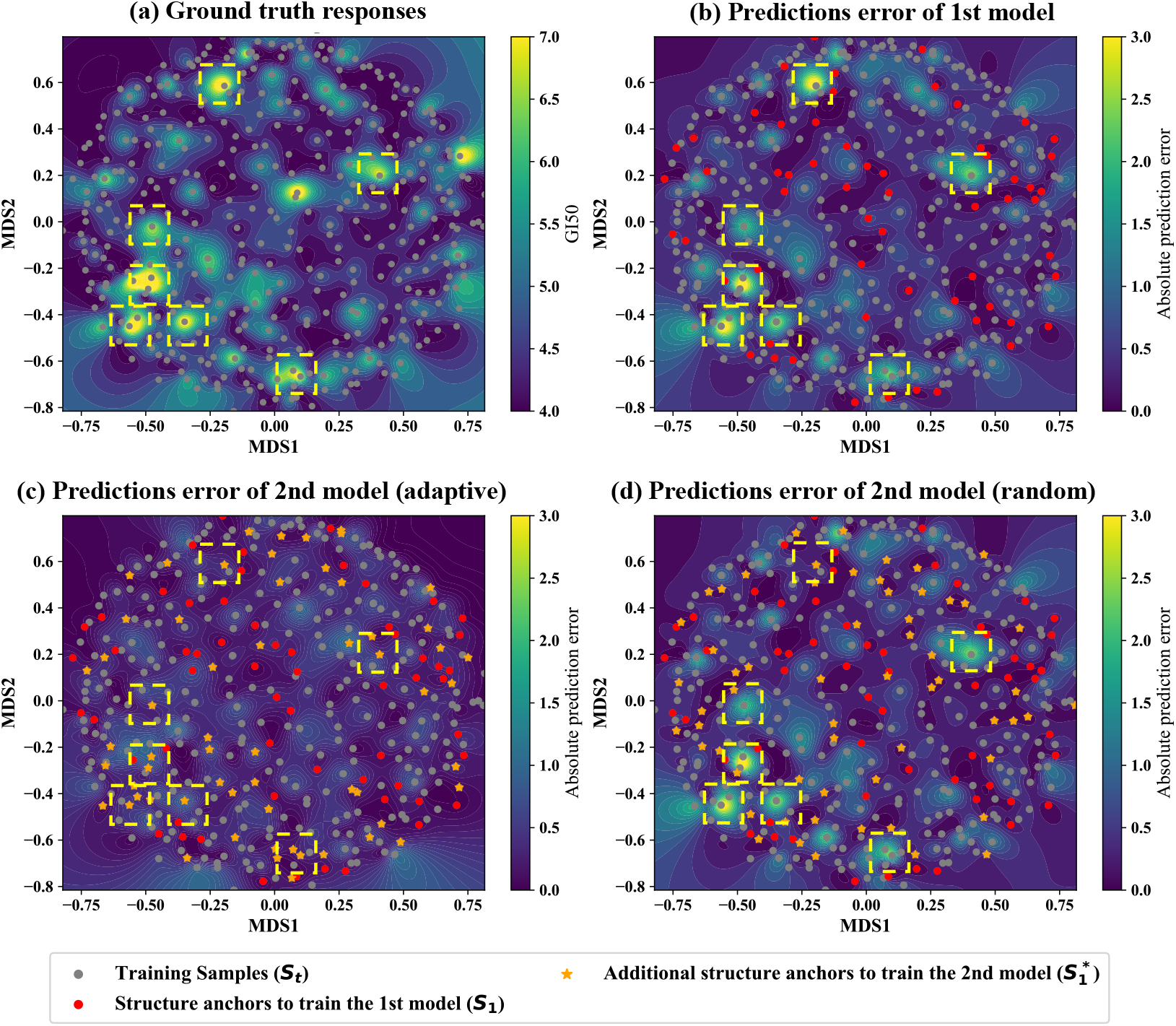
Comparison of absolute prediction error for TR models trained with random and adaptive anchor selection on 500 random samples from the HCC2998 cell line. The samples were projected onto a 2D structural space using multidimensional scaling (MDS), with landscapes interpolated using a Gaussian kernel. (a) Ground truth response values. The gray dots represent the samples. (b-c) Absolute prediction error for the (b) first and (c) second models in adaptive anchor selection. The red dots represent the 75 (15%) randomly selected structure anchors used to train the first model, denoted as ***S***_1_. The orange stars in (c) indicate the 75 samples that are not in *S*_1_ and showed the largest absolute prediction error from the first model, denoted as 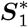. The combination of the red dots and the orange star was used to train the second model. (d)Absolute prediction error for the second model when 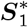 was randomly selected from samples not in ***S***_1_. The yellow boxes highlight the regions where the absolute prediction error from the first model is high.

In Figs. 4 (a-d), we used yellow boxes to highlight the regions where the absolute prediction error from the first model is high. In Fig. 4 (c), after selecting samples with high prediction error as structure anchors, the magnitude of the prediction error in the yellow-boxed regions is significantly reduced. In contrast, in Fig. 4 (d), the absolute prediction error is reduced in only one out of the seven yellow boxes because a sample with high prediction error in that box was randomly selected as a structure anchor. The prediction error of the other six boxes remains high since no high-error samples were picked as structure anchors in those regions. These results demonstrate that adaptive anchor selection strategically selects samples in regions where the current model performs poorly as structure anchors to effectively improve the performance of subsequent models.

#### 2.3.3 Comparing RBF-based and optimization-based response reconstruction

As mentioned in Section 1, the RBF-based reconstruction (see Eq. 5) fails to utilize large distance estimations, whereas the optimization-based reconstruction (see Eq. 9) ensures that all distance estimations contribute to the final prediction. In this section, we illustrate the distinction with simple toy examples shown in Fig. 5.

**Fig. 5:**
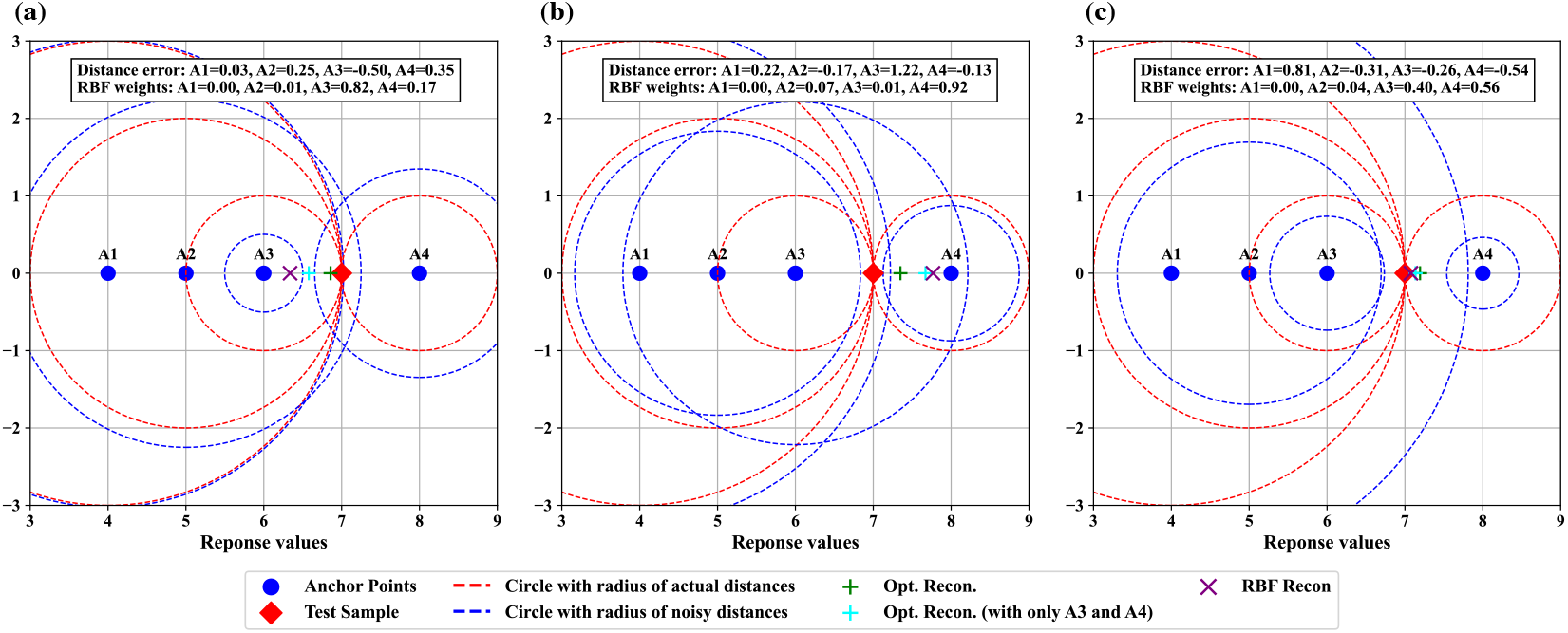
Toy examples comparing optimization-based and RBF-based response reconstruction with random Gaussian distance errors. In these examples, there are four anchor points (blue-filled circles) with response values of 4, 5, 6, and 8 (denoted as A1, A2, A3, and A4, respectively), and one test sample (red diamond) with a response value of 7. The red dashed circles represent the actual distances between the anchor points and the test sample. Random Gaussian noise with zero mean and a std of 0.5 is added to the actual distances. The noisy distances are represented by the blue dashed circles and are used to reconstruct the response of the test sample. The plus symbols indicate the responses reconstructed by the optimization-based approach with all anchors (green) and with only A3 and A4 (cyan). The purple cross symbol shows the response value reconstructed by the RBF-based approach. The text block in each figure lists the distance errors (i.e., the added Gaussian noise) and the weights used in RBF-based reconstruction. (a), (b), and (c) illustrate three cases where the distance errors are different.

In these examples, there are four response anchor points (blue-filled circles) with response values of 4, 5, 6, and 8 (denoted as A1, A2, A3, and A4, respectively), and one test sample (red diamond) with a response value of 7. As shown in Figs. S1, the response distance estimation error of the test samples from three representative cell lines has mean and std values around 0 and 0.5, respectively, and follows bell-shaped distributions that resemble a Gaussian distribution. Additionally, the magnitude of the estimation error is not linearly related to the actual response distances, as indicated by the PCC values being close to zero. Hence, in the toy examples, we added random Gaussian noise (mean=0, std=0.5) to the response distances between the test sample and anchor points to simulate response distance estimation error. The resulting noisy distances serve as simulated distance estimations and are used to reconstruct the response of the test sample using RBF-based and optimization-based approaches. Figs. 5 (a-c) show three cases where the Gaussian noise has different patterns.

For RBF-based reconstruction, in all three cases, the RBF weights assigned to A1 and A2 (the two anchors that are distant from the test sample) are close to zero, contributing negligibly to the final prediction. Consequently, the accuracy of the RBF-based approach depends primarily on the distance estimations of A3 and A4. In Fig. 5 (c), when the distance errors of A3 and A4 have the same sign and similar magnitude, the reconstructed response (purple cross) is close to the actual response. However, when the distance errors of A3 and A4 have opposite signs or different magnitudes, as shown in Figs. 5 (a) and (b), the reconstructed responses deviate significantly from the actual response.

In contrast, the responses reconstructed by the optimization-based approach (green plus) stay close to the actual response across all three cases. Furthermore, when A1 and A2 are excluded (because RBF assigns negligible weights to these samples) in the optimization-based reconstruction, the reconstructed responses (cyan plus) shift away from the actual response in Fig. 5 (a) and (b). These results demonstrate that large distance estimations contribute meaningfully to response reconstruction in the optimization-based approach.

### 2.4 Interpreting AdapToR model

In this section, we present an example of interpreting the AdapToR model trained on the HCC2998 cell line. By analyzing the model weights, we identified several key drugs and investigated how they impact AdapToR’s final prediction. The estimated weights, ***Ŵ***, is a *K*_*s*_ by *K*_*r*_ matrix, where *K*_*s*_ and *K*_*r*_ are the numbers of structure and response anchors. The estimated response distance matrix, 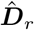, is calculated as 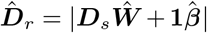, where ***D***_*s*_ is the structure distance matrix. The *i*th column of ***Ŵ*** is responsible for estimating the *i*th column in 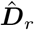 that corresponds to the distances associated with the *i*th response anchor. Thus, we define the *i*th column of ***Ŵ*** as the weights associated with the *i*th response anchor. Similarly, we define the *j*th row of ***Ŵ*** as the weights associated with the *j*th structure anchor. A positive (or negative) (*i, j*)-entry of ***Ŵ*** indicates a positive (or negative) relation between a sample’s structure distance to the *j*th structure anchor and its response distance to the *i*th response anchor.

Figs. 6 (a-b) show the top five structure anchors with the most positive (red) and negative (blue) weights associated with two representative response anchors: response anchor (A) (NSC# 638495 with NLOGGI50=4.28), and response anchor (B) (NSC# 699490 with NLOGGI50=7.86). Note that the NLOGGI50 values of all drugs range approximately from 4 to 8. For response anchor (A), a small structure distance to structure anchors with large positive weights would result in a small response distance to response anchor (A), suggesting a low response value. This aligns with our observation that the top five structure anchors with positive weights all exhibit low response values. In contrast, a small structure distance to structure anchors with large negative weights would lead to a large response distance to response anchor (A), indicating a high response value. As expected, all the top five structure anchors with negative weights show high response values. For response anchor (B), which has a high response value, we observe a reverse pattern: the top structure anchors with positive weights exhibit high response values, whereas those with large negative weights have low response values.

**Fig. 6:**
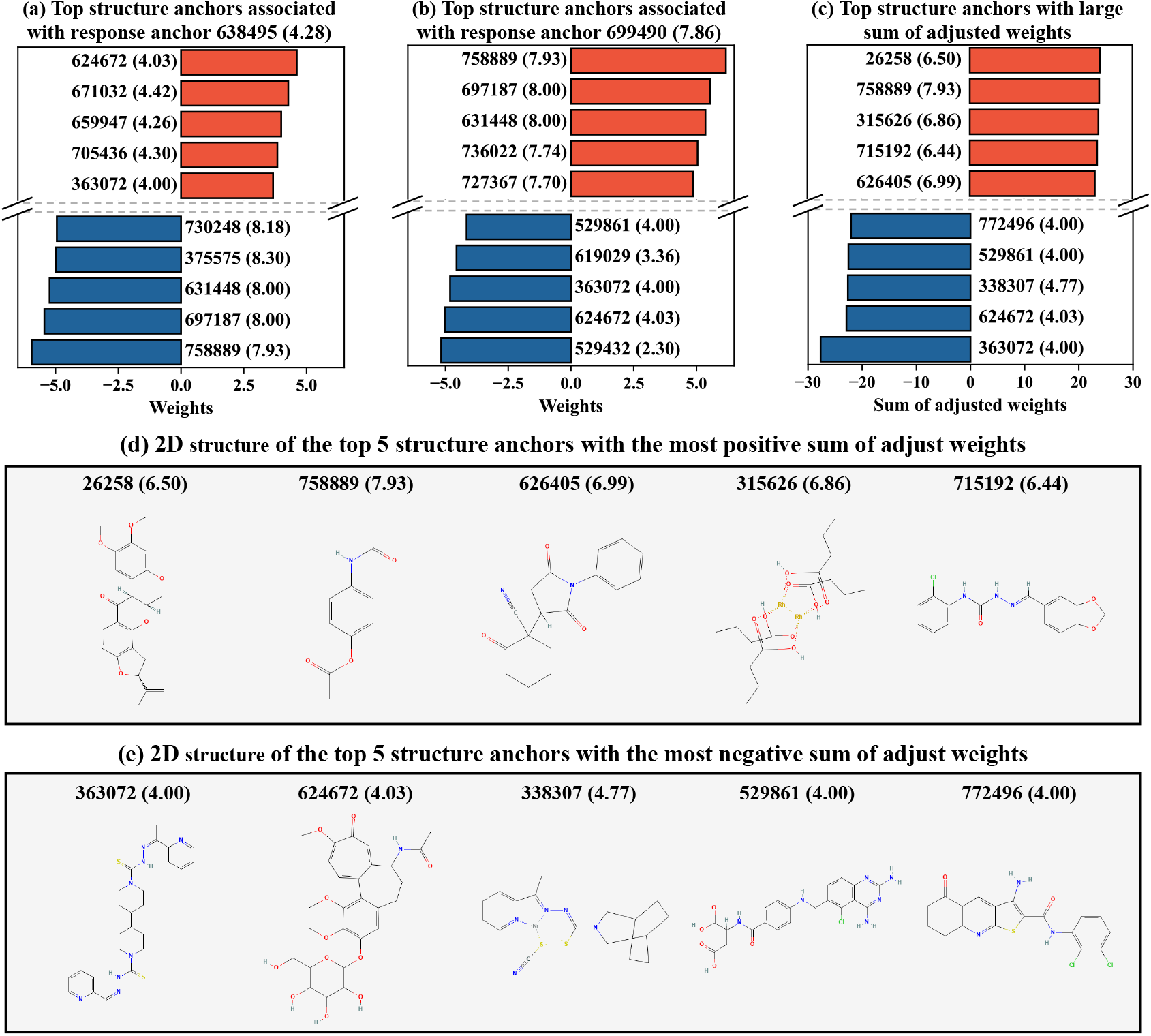
Interpreting the AdapToR model trained on the cell line HCC2998. (a-b) Top 5 structure anchors with the most positive (red) and negative (blue) weights associated with twp response anchors: (a) NSC# 638495 with NLOGGI50=4.28 and (b) NSC# 699490 with NLOGGI50=7.86. (c) Top 5 structure anchors with the most positive (red) and negative (blue) sum of adjusted weights over the response anchors. We adjusted ***Ŵ*** by multiplying -1 to the columns corresponding to the response anchors with response values smaller than 5 and summed up the adjusted weights along the columns. In (a-c), the drugs are represented as NSC# (NLOGGI50). (d-e) 2D structure of the top 5 structure anchors with the most (d) positive and (e) negative sum of adjusted weights.

**Fig. 7:**
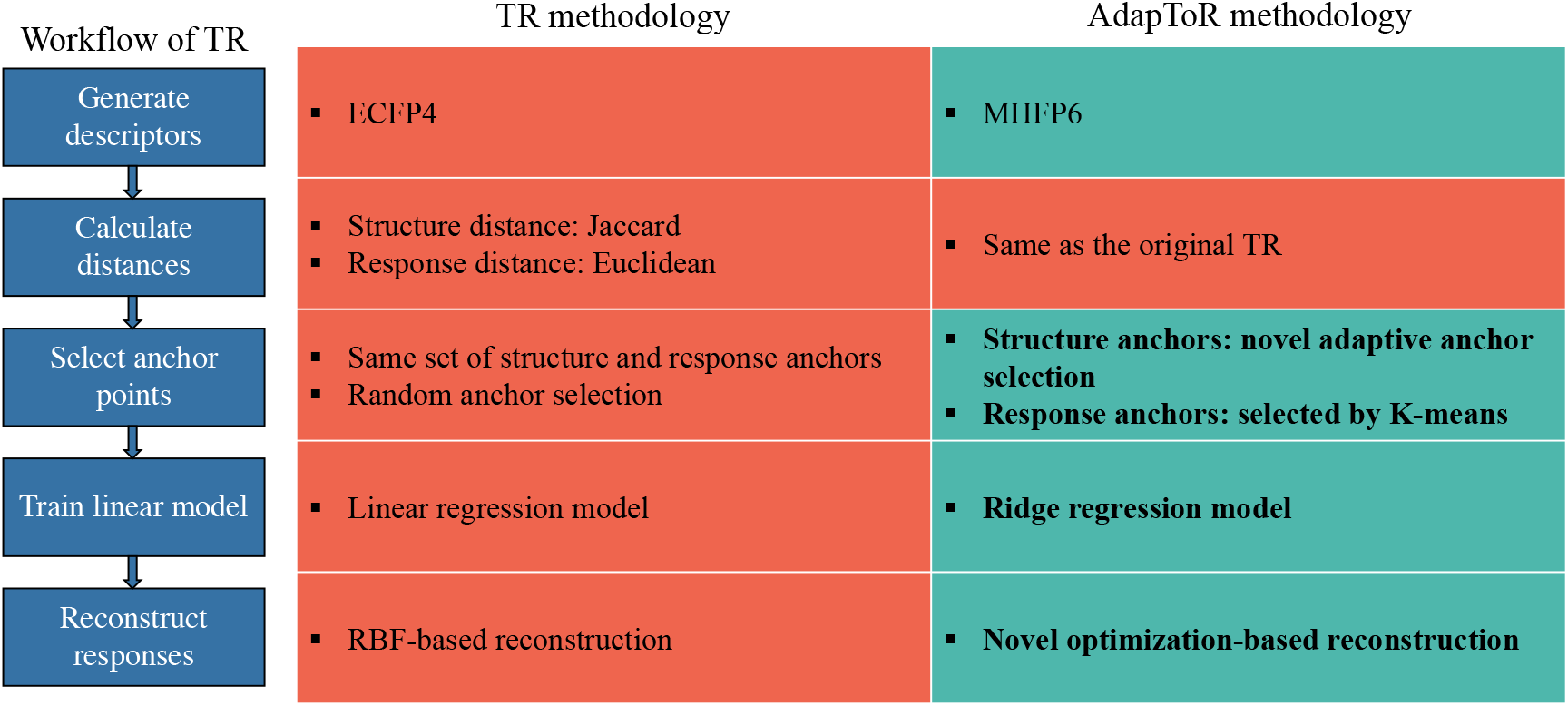
Comparison of TR and AdapToR methodologies. Green patches indicate modifications in AdapToR, while the red patch represents the TR methodology. Key features added in AdapToR are emphasized in bold.

Interestingly, we find common structure anchors in Figs. 6 (a) and (b). For example, structure anchor NSC# 798889 (with NLOGGI50=7.93) shows the most positive weight with response anchor (B) and the most negative weight with response anchor (A). While these weights have opposite signs, they deliver the same message that a small structure distance to NSC# 798889 would lead to a high response value. In contrast, structure anchor NSC# 624672 (with NLOGGI50= 4.03) exhibits a large negative weight with response anchor (B) and a large positive weight with response anchor (A), indicating that a large structure distance to NSC# 624672 would result in a high response value.

These findings motivate us to combine the weights across the response anchors to identify critical structure anchors for the HCC2998 cell line. To achieve this, we adjusted ***Ŵ*** by multiplying -1 to the columns corresponding to the response anchors with response values below 5 (an arbitrary threshold for low responses) and then summed the adjusted weights over the response anchors (i.e., along the columns). Structure anchors with large (positive or negative) sums of adjusted weights are considered to be critical, as a large response value can be associated with small (or large) structure distances to anchors with large positive (or negative) sums.

Fig. 6 (c) shows the top five structure anchors with the most positive (red) and negative (blue) sums of adjusted weights. The top five structure anchors showing the most positive sum of adjusted weights are NSC# 26258, 758889, 315626, 715192, and 626405, with their respective NLOGGI50 values of 6.50, 7.93, 6.86, 6.44, and 6.99. Their 2D structures are shown in Fig. 6 (d). The top five structure anchors with the most negative sum of adjusted weights are NSC# 363072, 624672, 338307, 529861, and 772496, with their respective NLOGGI50’s given by 4.00, 4.03, 4.77, 4.00, and 4.00. Their 2D structures are shown in Fig. 6 (e).

## 3 Discussion

In this work, we introduced **Adap**tive **To**pological **R**egression (AdapToR) for QSAR modeling. AdapToR builds upon Topological Regression (TR) [23] by incorporating features that enhance scalability, stability, interpretability, and predictive capacity. When tested on the NCI60 GI50 dataset, AdapToR demonstrates 5% to 17% improvements in performance metrics and significant reductions in training and testing times compared to TR. Also, AdapToR is compared against three baseline models: Linear Regression (LR), Random Forest (RF), and *K*− Nearest Neighbors (KNN), and two state-of-the-art deep learning models: Transformer-Convolutional Neural Network (TCNN) [12] and Graph Transformer (GraTrans) [17]. Our results show that AdapToR outperforms all competing models in all performance metrics. While TCNN with data augmentation (TCNN-Aug) produced predictive performance similar to AdapToR, our posited model achieved an order of magnitude gain in training and testing times. These results establish AdapToR as a powerful and computationally efficient tool for QSAR modeling.

In addition to the NCI60 dataset, we trained and evaluated AdapToR on 530 ChEMBL datasets. Again, AdapToR achieved the best predictive performance at considerably lower training and testing times compared to TCNN-Aug, demonstrating AdapToR’s superior performance across different datasets.

Another advantage of AdapToR is its interpretability. We provided an illustrative example demonstrating how a trained AdapToR model can be interpreted and how meaningful insights can be derived from it. By analyzing its weights across response anchors, we identified critical structure anchors shown in Figs. 6 (d) and (e). Our analyses suggest that a large response value can be associated with small structure distances to the drugs shown in (d) and large structure distances to the drugs shown in (e). We believe that revealing such associations can be helpful for hits-to-lead and lead optimization.

AdapToR incorporates four model features that enhance the stability, scalability, interpretability, and predictive capacity of the TR framework. First, the adoption of Ridge regression effectively addresses multicollinearity, an issue increasingly prevalent with larger datasets, thereby numerically stabilizing model behavior and significantly reducing training times. Second, AdapToR employs a reduced set of response anchors selected via *k*-means clustering to reduce model complexity while still ensuring adequate coverage of the response space. This clustering-based selection not only improves scalability but also enhances predictive performance by excluding redundant response anchors.

Furthermore, AdapToR includes two novel model features: adaptive structure anchor selection and optimization-based response reconstruction. The adaptive anchor selection approach strategically picks samples in underperformed areas as structure anchors. Compared to random anchor selection, adaptive selection results in structure anchors that are more informative for response prediction, hence enhancing model interpretability and predictive performance. Lastly, AdapToR reconstructs the optimal response under the loss objective stated in Eq. 9. This optimization-based approach is able to utilize all response distance estimates to improve the model’s predictive capacity. In addition, it leads to a roughly 10% decrease in testing time. While seemingly modest for thousands of test samples, this improvement can yield considerable computational savings in large-scale applications, such as virtual screening involving millions of compounds.

In conclusion, AdapToR is a powerful, reliable, computationally efficient, and interpretable tool for QSAR modeling. We anticipate that its implementation in real-world drug discovery and other QSAR tasks will further demonstrate its practical value and effectiveness.

## 4 Methods

### 4.1 Topological regression

Topological regression (TR) is a computationally efficient and highly interpretable QASR model recently proposed by [23]. TR builds linear models that use distances in the structure (input) space to predict distances in the response (output) space. The driving regression model in TR can be written as

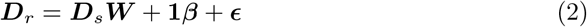

where 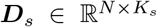 and 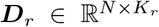 are the structure and response distance matrices, respectively, *N* is the number of samples, *K*_*s*_ and *K*_*r*_ are the numbers of structure and response anchors, respectively, 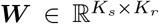 contains the model weights, **1** ∈ℝ^*N ×*1^ is an all-one vector, 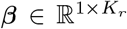 represents the intercept terms and 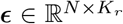 is the error term. In TR, 60% of the training samples are randomly selected as both structure and response anchors, resulting in *K*_*s*_ = *K*_*r*_ = round(0.6*N*). By defining 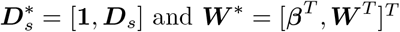, the equation can be rewritten as:

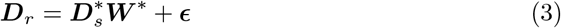

The least-square error solution ***Ŵ*** and the predicted response distance 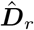 are given by:

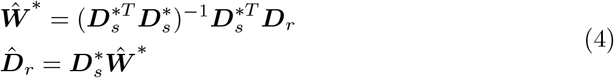

The response values are then reconstructed as follows:

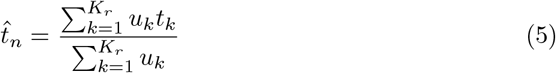

where 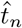 is the reconstructed response value for the *n*th sample, *t*_*k*_ is the response value of the *k*th response anchor. The predicted distances 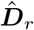 are converted to weights, *u*_*k*_, *k* = 1, 2, …, *K*_*r*_, using a Radial Basis Function (RBF) [23].

### 4.2 Adaptive Topological Regression (AdapToR)

In this work, we modified the TR methodology to enhance its scalability, stability, interpretability, and predictive capacity. Fig. 7 outlines the TR workflow and highlights the key model features added in AdapToR. These additional features include (1) the adoption of Ridge regression, (2) response anchor selection with *k*-means clustering, (3) adaptive structure anchor selection, and (4) optimization-based response reconstruction. The following sections provide detailed descriptions of these features. The pseudo-code for AdapToR is presented in Algorithm 1.

#### 4.2.1 Model features for enhancing scalability, stability, interpretability, and predictive capacity

##### Ridge regression

As demonstrated in Fig. S2, we observed that the condition number of the input distance matrix ***D***_*s*_ is exceptionally large, resulting in unstable model weights and abnormally long training times (see Table 2). To address this, we replaced the linear regression model with a Ridge regression model. Ridge regression adds an L2 penalty to the normal linear regression objective function as follows

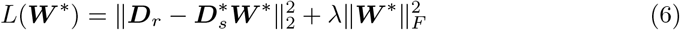

where ∥ · ∥_*F*_ calculates the Frobenius norm, which sums the squares of all elements in the matrix, and *λ* is a hyperparameter controlling the strength of the L2 penalty. In this work, we found that the optimal value for *λ* is 0.05 (see Section 4.2.3 for more details). By minimizing the objective function, the closed-form solution for Ridge regression is

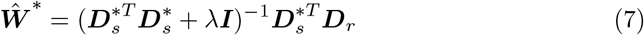

Since 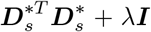 is well-conditioned (because of *λ****I***), we observed stable model weights (in Fig. S2) and reduced training time (in Table 2) after adding the L2-penalty.

##### Select response anchors with *k*-means clustering

To reduce model complexity, we used an independent and much smaller set of anchor points in the response space, making *K*_*r*_ *<< K*_*s*_. These response anchors were not selected randomly but were chosen by *k*-means clustering. Specifically, we applied *k*-means clustering in the response space with *K*_*r*_ clusters and selected the samples closest to the cluster centers as the response anchors. In this work, the optimal value for *K*_*r*_ was determined to be 10 (see Section 4.2.3 for more details). In Fig. S3, we presented the response values of the 10 anchors selected by *k*-means clustering in the response space for three representative cell lines. It can be observed that these anchors were distributed across the response space.

##### Adaptive anchor selection

As discussed in Section 1, TR randomly selects anchor points, resulting in uncertainty in model performance. Although the average of multiple independent TR models (i.e. Ensemble TR) can address such uncertainty, it reduces computational efficiency and model interpretability. To avoid this tradeoff, we proposed a novel adaptive anchor selection strategy that can preserve model interpretability while achieving better performance and lower computational costs compared to Ensemble TR. As shown in Algorithm 1, the adaptive anchor selection starts with randomly selecting *K*_*a*_ training samples as the initial set of structure anchors, denoted as ***S***_1_, to train the first TR model. Next, the training samples not included in ***S***_1_ are evaluated on the first model, and the top *K*_*a*_ samples with the highest absolute prediction error, denoted as 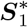, are added to the structure anchors to train the second model (i.e. 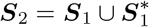). This process is repeated iteratively, with the *p*th model being trained on the set of structure anchors ***S***_*p*_, which is expressed as

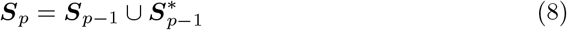

where ***S***_*p* − 1_ represents the set of structure anchors to train the (*p*− 1)th model and 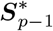 consists of the *K*_*a*_ training samples not in ***S***_*p* − 1_ that exhibit the highest absolute prediction error on the (*p* − 1)th model. After completing the iterative process, the last model is used for evaluation. The adaptive anchor selection has two hyperparameters: the number of structure anchors selected at each step *K*_*a*_ and the number of steps (i.e., models) *P* . Section 4.2.3 describes how these hyperparameters are fine-tuned.

##### Optimization-based response reconstruction

With the predicted distances between the samples and the response anchors, TR reconstructs the responses as shown in Eq. 5. As mentioned in Section 1, the RBF-based reconstruction fails to utilize large response distance estimates and is not optimized. Therefore, we proposed an optimization-based reconstruction method that can utilize information across all distance estimates and is optimized under the stated loss criterion. The response of the *n*th sample (denoted as 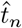) is determined by minimizing the sum of the square of the differences between *max*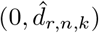 and 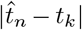 across all response anchors, which can be expressed as

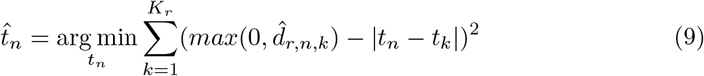

where 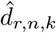 is the estimated response distance between the *n*th sample and *k*th response anchor and *t*_*k*_ is the response value of the *k*th response anchor. The function *max*(·,·) ensures that the distance estimations are non-negative. We used the Nelder–Mead algorithm to solve the optimization problem [29].

#### 4.2.2 Statistical Analysis of AdapToR

Our goal in this section is to offer a statistical formulation to understand the statistical coherence of the AdapToR technique. [23] showed that the original TR admits formal statistical inference. Since we had modified the prediction strategy in AdapToR (using Eq. 9 instead of Eq. 5), we theoretically show that AdapToR also admits a coherent Bayesian hierarchical model and, therefore, is amenable to formal statistical inference.

Observe, if we ignore the cross-product term obtained after expanding Eq. 9, then 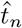 is simply the mean of *t*_1_, *t*_2_, …, *t*_*K*_ with 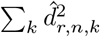 providing the offset term (because at the time of response reconstruction, 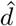 is already known). The cross-product term 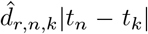 forces the distance in the response space, after reconstruction, to be positively associated with estimated distances and hence ensures convexity of Eq. 9.

So, from inferential perspective, we treat response reconstruction as an estimation problem. We mimic the cross-product term of Eq. 9 to construct a concave joint conditional distribution of 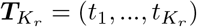 as follows:

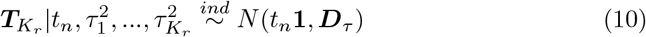

where 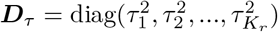. Next, we define conditionally independent exponential distribution for each 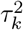 as follows:

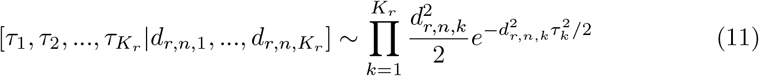

##### Result

For each *k* = 1, 2, …, *K*_*r*_ the distribution of *t*_*k*_| *t*_*n*_, *d*_*r*,*n*,*k*_ is *Laplace*(*t*_*n*_, 1*/d*_*r*,*n*,*k*_)

*Proof:* For each *k*, conditional moment-generating function (MGF) of *t*_*k*_

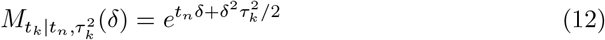

Integrating over *τ*_*k*_, the resulting MGF is given by

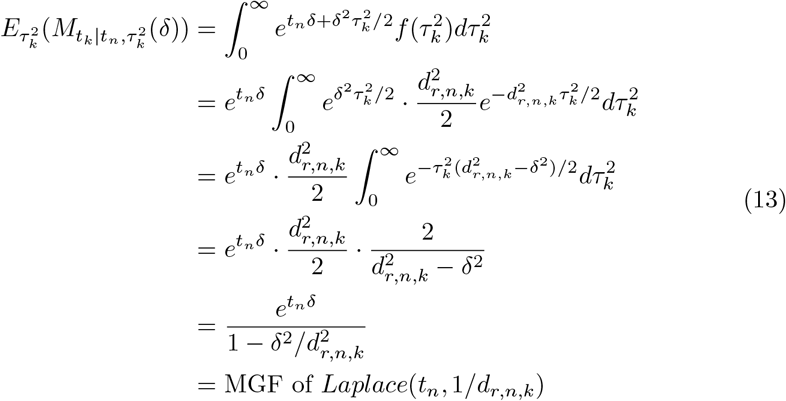

Thus, after integrating out 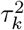, we have

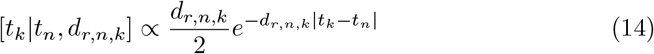

Observe that if we exponentiate the cross product term in Eq.9, it yields 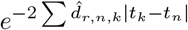, which is essentially the same as the exponential term of the joint distribution of *t*_1_, *t*_2_, …, *t*_*K*_ *t*_*n*_, *d*_*r*,*n*,1_, …, *d*_*r*,*n*,*K*_ .

So, the optimization problem in Eq.9 can be treated as a statistical point estimation problem under the following hierarchy

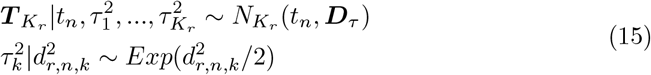

where 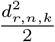 is the rate parameter.

At the so-called process level, we have:

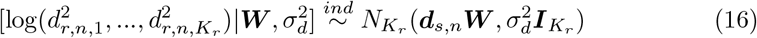

where ***d***_*s*,*n*_ is the *n*th row of ***D***_*n*_ in Eq.2. Note that, under this construction, we only need ***D***_*r*_ to contain non-negative elements. They need not be proper metrics. Consequently, we do not need to impose positive semi-definiteness when predicting 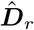, we are essentially predicting the mean of the precision term associated with the model for 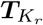 .

To accommodate Ridge regularization, following [30], we impose the following priors on the elements of ***W***

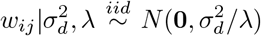

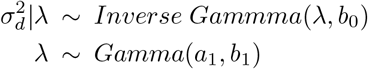

Note that, the above *λ* is the ridge parameter. But, under Bayes paradigm, it is a random variable admitting a distribution on positive support. To complete the hierarchy, we specify a Normal(0, *σ*^2^) prior on *t*_*n*_.

##### Result

Under the above hierarchical model, the conditional specification of the response variable is preserved. In other words, the conditional distribution of 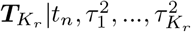 belongs to the same family of distributions as the full conditional posterior distribution of 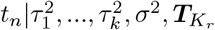 .

*Proof*. We only need to prove that the full conditional of 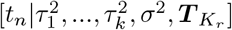 is Normal.

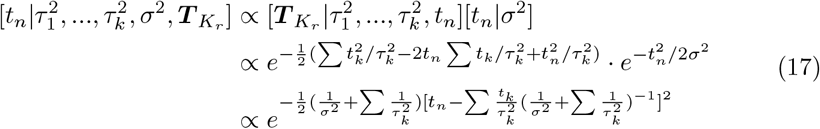

Thus, 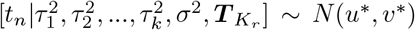 where 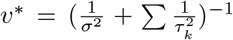 and 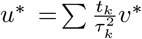.

Thus, the conditional posterior distribution of *t*_*n*_ is unimodal and *u*^*∗*^ is the point estimate of *t*_*n*_. We can explicitly see how the values of the response anchors determine the estimates of reconstructed responses. The estimated values of 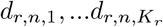 enter into the point estimate of *t*_*n*_ via 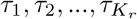 .

#### 4.2.3 Hyperparameters fine-tuning

In AdapToR, we have four hyperparameters: (1) *λ*, which controls the L2 penalty; (2) the number of response anchors, *K*_*r*_; (3) the number of structure anchors selected at each step of adaptive anchor selection, *K*_*a*_; and (4) the number of steps, *P* . We utilized the Python package *Optuna* with the tree-structure Parzen estimator [31] to fine-tune the hyperparameters on the training samples of three representative cell lines. For each cell line, the training samples were divided into training (80%), testing (10%), and validation (10%). Note that these splits were solely for hyperparameter tuning.

The optimal values for the hyperparameters were determined by minimizing the NRMSE evaluated on the test samples over 100 trials. Fine-tuning *λ* and *K*_*r*_ is straightforward. However, it may not be optimal or feasible to directly fine-tune *K*_*a*_ and *P* . For *K*_*a*_, we fine-tuned the percentage of structure anchors, calculated as 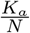, which can be more generalizable to cell lines with different numbers of drugs. For *P*, directly fine-tuning it may cause potential issues as its feasible range depends on the value of 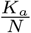. Alternatively, we implemented early-stopping, monitored by validation samples, and fine-tuned the early-stopping hyperparameter *min delta* (with a fixed patience of 2). The number of steps used with the optimal *min delta* was regarded as the optimal value for *P* .

Table 4 summarizes the suggested value ranges and fine-tuned values for the hyperparameters. In this work, the hyperparameters were set to *λ* = 0.05, *K*_*a*_ = 10, 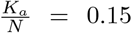 and *P* = 4, based on the averaged optimal values over the three representative cell lines.

**Table 4:** Fine-tuned hyperparameters with three representative cell lines: HCC2998, 786-O and Colo205

| Hyperparameter | Suggested range | HCC2998 | 786-O | Colo205 | Average | Value used |
| --- | --- | --- | --- | --- | --- | --- |
| $\lambda$ | 0.01-0.2 | 0.05 | 0.1 | 0.03 | 0.06 | 0.05 |
| $K_a$ | 5-15 | 14 | 10 | 10 | 11 | 10 |
| $\frac{K_a}{N}$ | 0.01-0.2 | 0.14 | 0.19 | 0.16 | 0.16 | 0.15 |
| $min\_delta$ | 0.001-0.01 | 0.004 | 0.005 | 0.009 | 0.006 | \ |
| $P$ | \ | 4 | 4 | 4 | 4 | 4 |

$\lambda$ : it controls L2-penalty, $K_r$ : # of response anchors, $\frac{K_a}{N}$ : percent of structure anchors added at each step of adaptive anchor selection, $P$ : # of steps in adaptive anchor selection.

### 4.3 QSAR models for comparison

#### 4.3.1 Baseline models

We implemented and evaluated several commonly used baseline QSAR models, including random forest (RF), linear regression (LR), LR with L1-penalty (LASSO), LR with L2-penalty (Ridge), and *K*− Nearest Neighbors (KNN) regression. These models take molecular fingerprints as input features to predict the responses. As shown in Fig. S4, all models showed the lowest NRMSE with ECFP4 as input. Also, we observed that LR and Ridge showed similar NRMSE and outperformed LASSO. Therefore, only the LR results are included in the main text. For KNN, the number of nearest neighbors was set to 5, as increasing the value to 10 or 15 led to higher NRMSE. RF was implemented with the *scikit-learn* function *RandomForestRegressor* with default settings.

#### 4.3.2 Transformer-Convolutional Neural Network

Transformer-Convolutional Neural Network (TCNN) is a powerful DL model, described as a Swiss-army knife for QSAR modeling [12]. The TCNN architecture combines a pre-trained Transformer encoder, trained on over 17 million string pairs for SMILES canonicalization, with a task-specific Text-CNN model for predictions. Additionally, TCNN supports data augmentation in testing and training by generating multiple non-canonical SMILES for each compound. We followed the TCNN instructions and trained the model on the SMILES strings, with and without augmentation, for up to 35 epochs using learning rate scheduling and early stopping. TCNN with augmentation is denoted as TCNN-Aug.

#### 4.3.3 Graph Transformer

Graph Transformer (GraTrans) is a DL model that utilizes graph representations of the drugs to predict drug responses [17]. The model architecture consists of Transformer encoders, Graph Attention Network (GAT) and Graph Convolutional Network (GCN) layers to extract features from molecular graphs, followed by two fully connected layers for response prediction (model architecture is shown in Fig. S5). The hyperparameters were set as follows: dropout rate = 0.5, learning rate = 10^*−*4^, and batch size = 512. These hyperparameters were fine-tuned using the Python package *Optuna* over 100 trials on the HCC2998 cell line. The model was trained for up to 200 epochs with learning rate scheduling and early stopping.

## 5 Acknowledgements

This work was supported in part by the National Science Foundation under Grants Nos. 2007903 (received by R.P.) and 2007418 (Received by S.G) and Leidos Biomed/NCI under contract 22X049. Any opinions, findings, and conclusions or recommendations expressed in this material are those of the authors and do not necessarily reflect the views of the National Science Foundation or Leidos Biomed/NCI.

## 6 Author contributions

Y.M., S.G., and R.P. formulated the problem and conceived the experiments, Y.M conducted the experiments and drafted the manuscript, Y.M, S.G, and R.P. analyzed the results, critically reviewed and edited the manuscript. All authors approved the final version.

## 7 Data and code availability

NCI60 GI50 dataset is available in the Development Therapeutic Program (DTP) repository (https://dtp.cancer.gov/databases_tools/bulkdata.htm). The ChEMBL datasets are available in the ChEMBL database (https://www.ebi.ac.uk/chembl/). AdapToR is publicly available at https://github.com/yixmao/adaptor.

## 9 Tables

### Algorithm 1

AdapToR

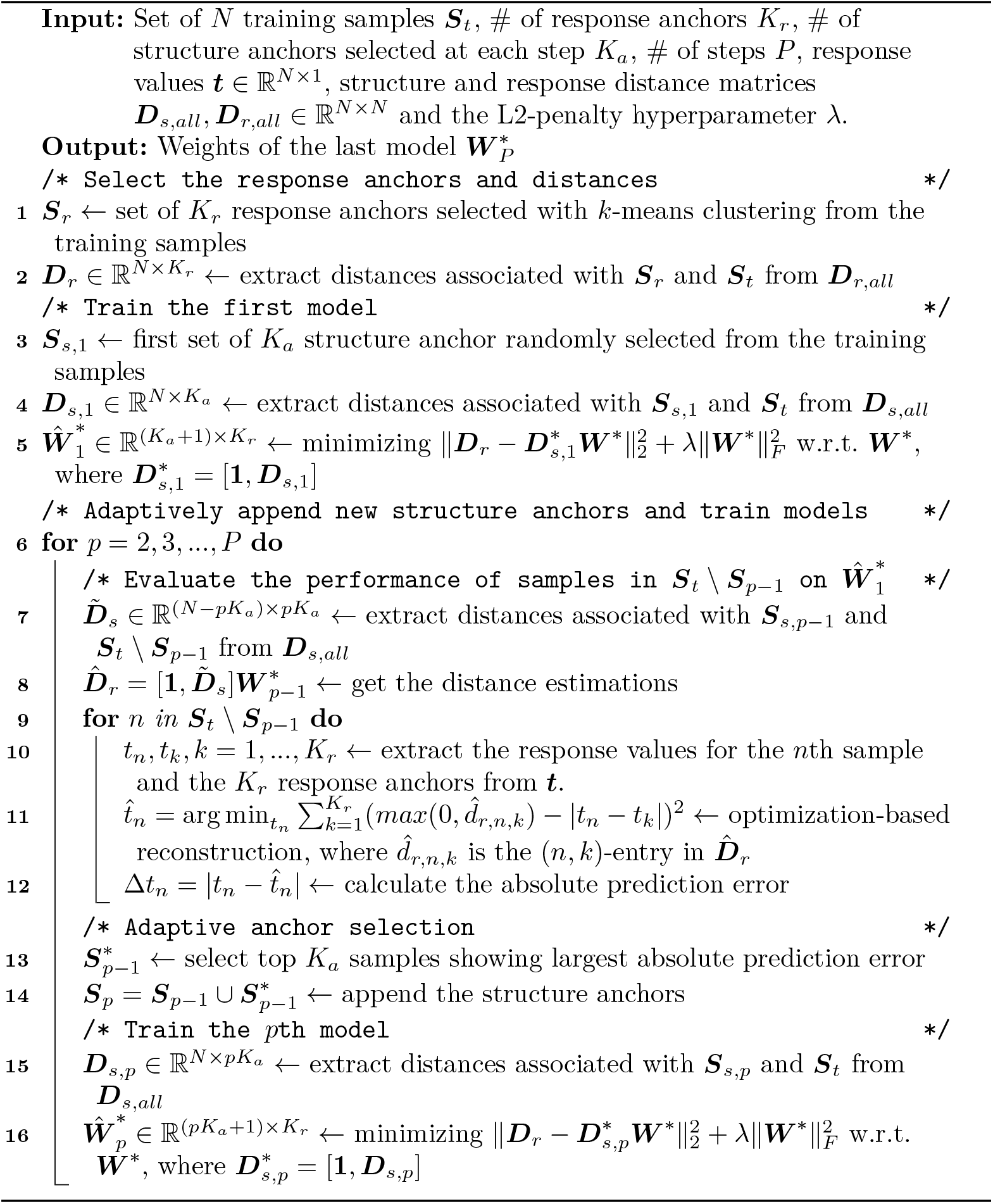

## References

[1] Neves, B. J. et al. QSAR-Based Virtual Screening: Advances and Applications in Drug Discovery. Frontiers in Pharmacology 9, 1275 (2018). doi:10.3389/fphar.2018.01275.

[2] Suay-Garcia, B. et al. Quantitative structure–activity relationship methods in the discovery and development of antibacterials. Wiley Interdisciplinary Reviews: Computational Molecular Science 10(6) (2020). doi:10.1002/wcms.1472.

[3] Cherkasov, A. et al. QSAR Modeling: Where Have You Been? Where Are You Going To? Journal of Medicinal Chemistry 57(12), 4977–5010 (2014). doi:10.1021/jm4004285.

[4] Perkins, R., Fang, H., Tong, W. & Welsh, W. J. Quantitative structure-activity relationship methods: Perspectives on drug discovery and toxicology. Environmental Toxicology and Chemistry 22(8), 1666–1679 (2003). doi:10.1897/01-171.

[5] Rogers, D. & Hahn, M. Extended-Connectivity Fingerprints. Journal of Chemical Information and Modeling 50(5), 742–754 (2010). doi:10.1021/ci100050t.

[6] Probst, D. & Reymond, J.-L. A probabilistic molecular fingerprint for big data settings. Journal of Cheminformatics 10(1), 66 (2018). doi:10.1186/s13321-018-0321-8.

[7] Moriwaki, H., Tian, Y.-S., Kawashita, N. & Takagi, T. Mordred: a molecular descriptor calculator. Journal of Cheminformatics 10(1), 4 (2018). doi:10.1186/s13321-018-0258-y.

[8] Svetnik, V. et al. Random Forest: A Classification and Regression Tool for Compound Classification and QSAR Modeling. Journal of Chemical Information and Computer Sciences 43(6), 1947–1958 (2003). doi:10.1021/ci034160g.

[9] Nagamine, N. & Sakakibara, Y. Statistical prediction of protein–chemical interactions based on chemical structure and mass spectrometry data. Bioinformatics 23(15), 2004–2012 (2007). doi:10.1093/bioinformatics/btm266.

[10] Robinson, R. L. M., Palczewska, A., Palczewski, J. & Kidley, N. Comparison of the Predictive Performance and Interpretability of Random Forest and Linear Models on Benchmark Data Sets. Journal of Chemical Information and Modeling 57(8), 1773–1792 (2017). doi:10.1021/acs.jcim.6b00753.

[11] Wang, S., Guo, Y., Wang, Y., Sun, H. & Huang, J. SMILES-BERT. Proceedings of the 10th ACM International Conference on Bioinformatics, Computational Biology and Health Informatics 429–436 (2019). doi:10.1145/3307339.3342186.

[12] Karpov, P., Godin, G. & Tetko, I. V. Transformer-CNN: Swiss knife for QSAR modeling and interpretation. Journal of Cheminformatics 12(1), 17 (2020). doi:10.1186/s13321-020-00423-w, 1911.06603.

[13] Nguyen, T., Nguyen, G. T. T., Nguyen, T. & Le, D.-H. Graph Convolutional Networks for Drug Response Prediction. IEEE/ACM Transactions on Computational Biology and Bioinformatics 19(1), 146–154 (2020). doi:10.1109/tcbb.2021.3060430.

[14] Bazgir, O. et al. Representation of features as images with neighborhood dependencies for compatibility with convolutional neural networks. Nature Communications 11(1), 4391 (2020). doi:10.1038/s41467-020-18197-y, 1912.05687.

[15] Wang, H. et al. GADRP: graph convolutional networks and autoencoders for cancer drug response prediction. Briefings in Bioinformatics 24(1), bbac501 (2022). doi:10.1093/bib/bbac501.

[16] Ma, T. et al. DualGCN: a dual graph convolutional network model to predict cancer drug response. BMC Bioinformatics 23(Suppl 4), 129 (2022). doi:10.1186/s12859-022-04664-4.

[17] Chu, T., Nguyen, T. T., Hai, B. D., Nguyen, Q. H. & Nguyen, T. Graph Transformer for Drug Response Prediction. IEEE/ACM Transactions on Computational Biology and Bioinformatics 20(2), 1065–1072 (2023). doi:10.1109/tcbb.2022.3206888.

[18] Rudin, C. Stop explaining black box machine learning models for high stakes decisions and use interpretable models instead. Nature Machine Intelligence 1(5), 206–215 (2019). doi:10.1038/s42256-019-0048-x.

[19] Bach, S. et al. On Pixel-Wise Explanations for Non-Linear Classifier Decisions by Layer-Wise Relevance Propagation. PLoS ONE 10(7), e0130140 (2015). doi:10.1371/journal.pone.0130140.

[20] Montavon, G., Binder, A., Lapuschkin, S., Samek, W. & Müller, K.-R. Layerwise relevance propagation: an overview. Explainable AI: interpreting, explaining and visualizing deep learning 193–209 (2019) .

[21] Linardatos, P., Papastefanopoulos, V. & Kotsiantis, S. Explainable AI: A Review of Machine Learning Interpretability Methods. Entropy 23(1), 18 (2020). doi:10.3390/e23010018.

[22] Yamanishi, Y., Araki, M., Gutteridge, A., Honda, W. & Kanehisa, M. Prediction of drug–target interaction networks from the integration of chemical and genomic spaces. Bioinformatics 24(13), i232–i240 (2008). doi:10.1093/bioinformatics/btn162.

[23] Zhang, R., Nolte, D., Sanchez-Villalobos, C., Ghosh, S. & Pal, R. Topological regression as an interpretable and efficient tool for quantitative structureactivity relationship modeling. Nature Communications 15(1), 5072 (2024). doi:10.1038/s41467-024-49372-0.

[24] Gaulton, A. et al. ChEMBL: a large-scale bioactivity database for drug discovery. Nucleic Acids Research 40(D1), D1100–D1107 (2012). doi:10.1093/nar/gkr777.

[25] Gaulton, A. et al. The ChEMBL database in 2017. Nucleic Acids Research 45(D1), D945–D954 (2017). doi:10.1093/nar/gkw1074.

[26] Yang, K. et al. Analyzing Learned Molecular Representations for Property Prediction. Journal of Chemical Information and Modeling 59(8), 3370–3388 (2019). doi:10.1021/acs.jcim.9b00237.

[27] Shoemaker, R. H. The NCI60 human tumour cell line anticancer drug screen. Nature Reviews Cancer 6(10), 813–823 (2006) .

[28] Rdkit: Open-source cheminformatics software. https://www.rdkit.org. Accessed: 2025-2-18.

[29] Gao, F. & Han, L. Implementing the Nelder-Mead simplex algorithm with adaptive parameters. Computational Optimization and Applications 51(1), 259–277 (2012). doi:10.1007/s10589-010-9329-3.

[30] Amin, M., Akram, M. N. & Ramzan, Q. Bayesian estimation of ridge parameter under different loss functions. Communications in Statistics - Theory and Methods 51(12), 4055–4071 (2022). doi:10.1080/03610926.2020.1809675.

[31] Watanabe, S. Tree-structured Parzen estimator: Understanding its algorithm components and their roles for better empirical performance (2023).

