## Supplementary Material for "AdapToR: Adaptive Topological Regression for quantitative structure-activity relationship modeling"

Table S1: Number of drugs for 60 cell lines in the NCI60 dataset

| Cell Line | # of drug[1] | Cell Line | # of drug[1] | Cell Line | # of drug[1] |
|---|---|---|---|---|---|
| 786-O | 48701 | K562 | 48264 | RXF 393 | 44464 |
| A498 | 43186 | LOX IMVI | 46786 | SF268 | 49153 |
| A549 | 49910 | M14 | 48659 | SF295 | 49356 |
| ACHN | 49002 | MALME-3M | 45808 | SF539 | 46680 |
| BT549 | 33183 | MCF7 | 36618 | SK-MEL-2 | 46433 |
| CAKI | 46706 | MDA-MB-231 | 36009 | SK-MEL-28 | 48801 |
| CCRF-CEM | 46820 | MDA-MB-435 | 36573 | SK-MEL-5 | 48560 |
| Colo205 | 49065 | MOLT-4 | 48608 | SK-OV-3 | 47124 |
| DU145 | 36321 | NCI-ADR-RES | 36855 | SN12C | 49218 |
| EKVX | 46990 | NCI-H226 | 45981 | SNB-19 | 49017 |
| HCC2998 | 44474 | NCI-H23 | 49310 | SNB-75 | 45957 |
| HCT-116 | 49184 | NCI-H322M | 48066 | SR | 40869 |
| HCT-15 | 49219 | NCI-H460 | 48230 | SW620 | 49784 |
| HL-60 (TB) | 44764 | NCI-H522 | 45347 | TK-10 | 48137 |
| HOP-62 | 48208 | OVCAR-3 | 48270 | T-47D | 34468 |
| HOP-92 | 43665 | OVCAR-4 | 47199 | U251 | 49466 |
| HS 578T | 34512 | OVCAR-5 | 48478 | UACC-257 | 48992 |
| HT29 | 49145 | OVCAR-8 | 49794 | UACC-62 | 48549 |
| IGROV1 | 48925 | PC-3 | 36407 | UO-31 | 48917 |
| KM12 | 49282 | RPMI-8226 | 46236 | **MDA-MB-468**[2] | 5566 |

[1] Drugs with invalid SMILES are excluded.
[2] Cell line MDA-MB-468 was excluded from the analysis due to a limited number of available drugs.

Table S2: Model performance averaged over 530 ChEMBL datasets and 5-fold cross-validation splits

| Model | NRMSE | Spearman | Train time (sec) | Test time (sec) |
|---|---|---|---|---|
| TCNN | 0.660 | 0.744 | 30.6 | 6.9 |
| TCNN-Aug | 0.596 | 0.786 | 109.8 | 20.0 |
| TR | 0.626 | 0.763 | **1.6** | 1.0 |
| Ensemble TR[1] | 0.599 | 0.785 | 13.8 | 12.3 |
| AdapToR | **0.586** | **0.791** | **1.7** | **0.2** |

[1] Ensemble TR in this table refers to Ensemble TR described in [2], which does not incorporate any new model features proposed in the main text.

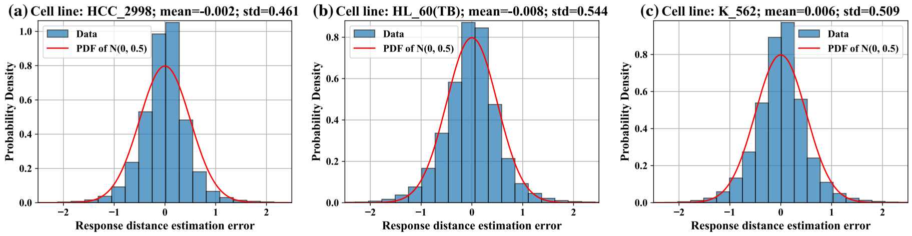


Figure S1: (a-c) Distributions of response distance estimation error of the test samples for three representative cell lines: (a) HCC2998, (b) HL-60 (TB), and (c) K562. As indicated in the titles, the mean and std values of the estimation error are around 0 and 0.5, respectively. Because the error distributions exhibit a bell-shaped pattern, we compared them with a Gaussian distribution (mean=0 and std=0.5), whose probability density function (PDF) is overlaid in each plot. It can be observed that the actual distributions are similar to the Gaussian distribution. To assess whether the magnitude of the estimation error correlates with actual response distances, Pearson's correlation coefficient (PCC) was computed. The PCC values are 0.086, 0.066, and 0.087 for cell lines HCC2998, HL-60 (TB), and K562, respectively.

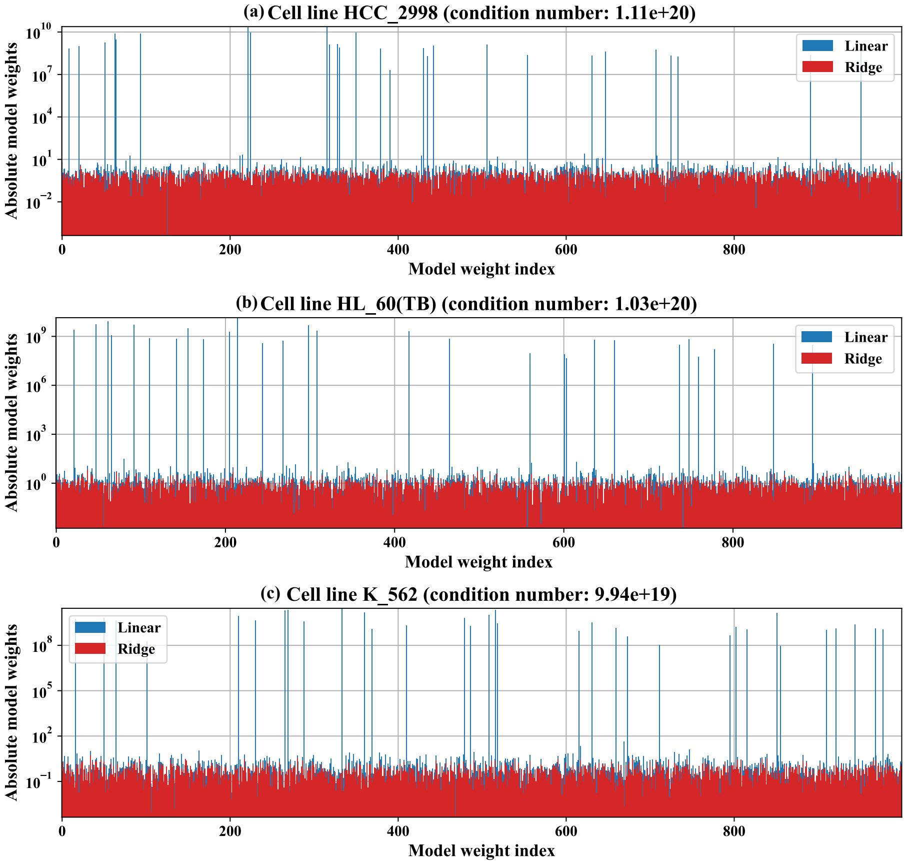


Figure S2: Absolute weights of linear TR models (blue) and Ridge TR models (red) trained on three representative cell lines: (a) HCC2998, (b) HL-60 (TB), and (c) K562. In each case, 60% of the training samples were selected as structure anchors, and the first 1,000 model weights were plotted. The figure titles indicate the condition number of the design matrix for each cell line. Notably, extraordinarily large condition numbers lead to unstable weights in the linear models, whereas the Ridge models have stable weights because of the L2 penalty.

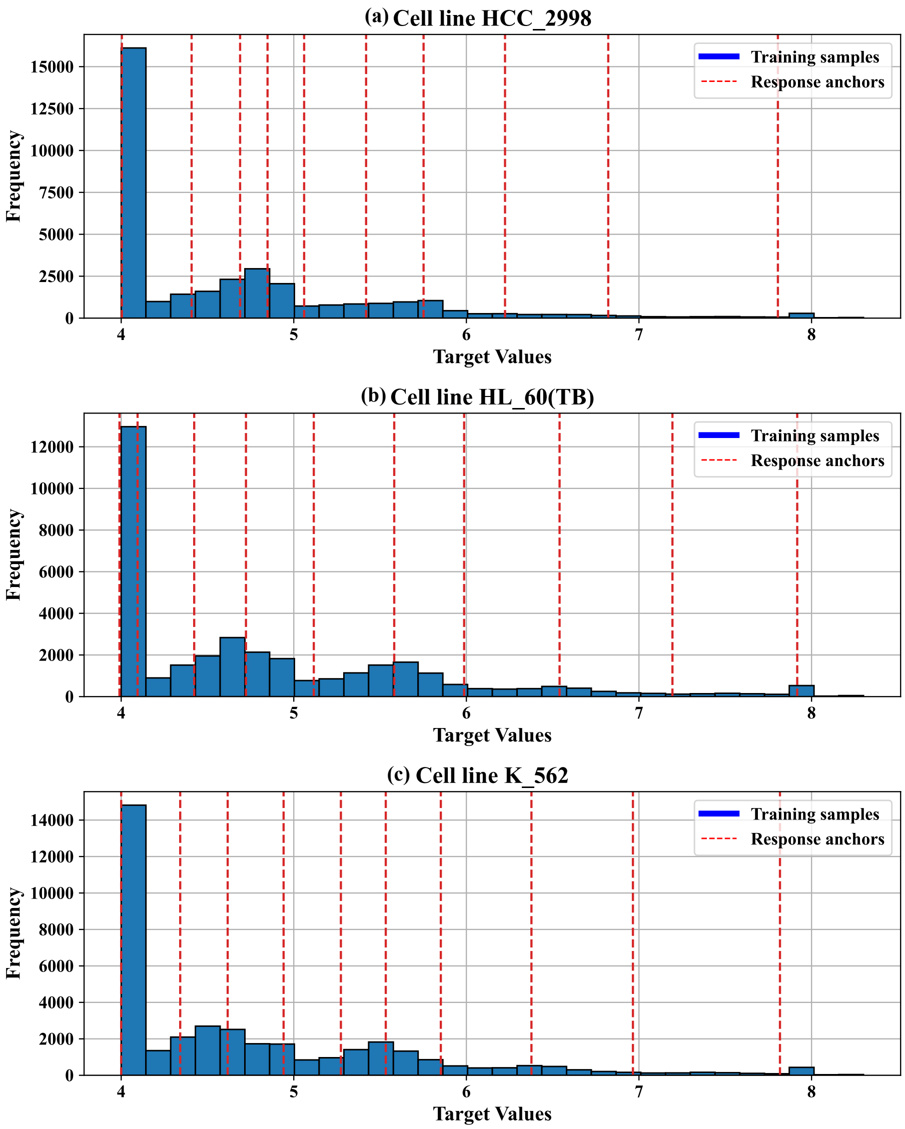


Figure S3: Histograms of response values for training samples for three representative cell lines: (a) HCC2998, (b) HL-60 (TB), and (c) K562. Dashed lines indicate response values of the 10 response anchors selected by K-means clustering.

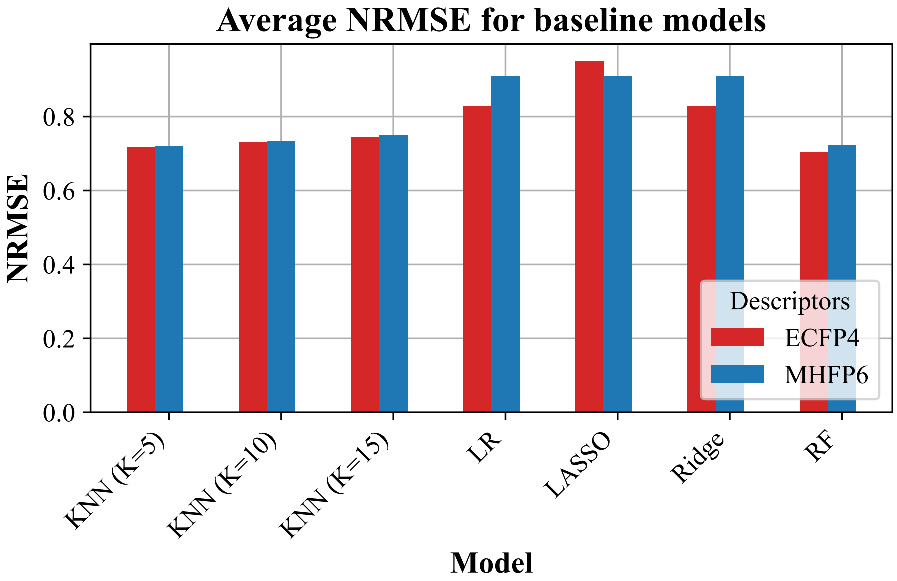


Figure S4: Average NRMSE of baseline models, including LR, LASSO, Ridge, RF, and KNN with ECFP4 and MHFP6 as input. The NRMSE values were averaged over 59 cell lines and 5 CV splits. For KNN, the number of nearest neighbors ($K$) was set to 5, 10, and 15. For LASSO and Ridge, the hyperparameter $\lambda$ was set to 0.05. LR: Linear Regression, LASSO: Linear regression with L1-regularization, Ridge: Linear regression with L2-regularization, RF: Random Forest, KNN: k-nearest neighbor.

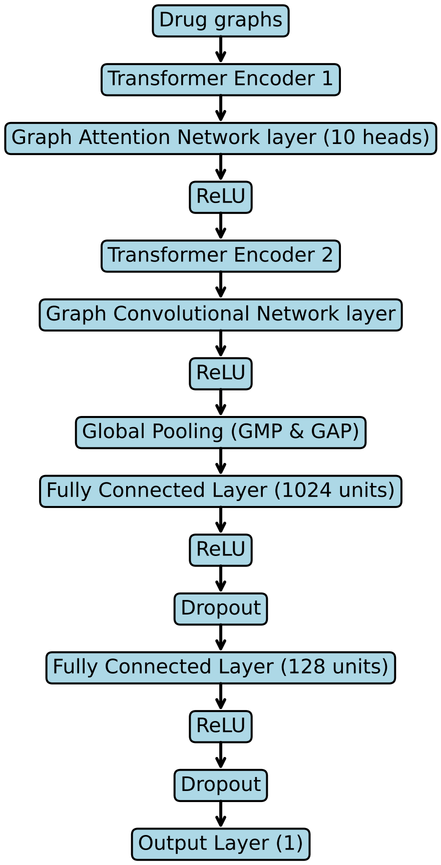


Figure S5: Model architecture of Graph Transformer adapted from [1].

### References


[1] T. Chu, T. T. Nguyen, B. D. Hai, Q. H. Nguyen, and T. Nguyen. Graph Transformer for Drug Response Prediction. *IEEE/ACM Transactions on Computational Biology and Bioinformatics*, 20(2):1065–1072, 2023.

[2] R. Zhang, D. Nolte, C. Sanchez-Villalobos, S. Ghosh, and R. Pal. Topological regression as an interpretable and efficient tool for quantitative structure-activity relationship modeling. *Nature Communications*, 15(1):5072, 2024.
